# Genome plasticity, a key factor of evolution in prokaryotes

**DOI:** 10.1101/357400

**Authors:** Itamar Sela, Yuri I. Wolf, Eugene V. Koonin

**Affiliations:** National Center for Biotechnology Information, National Library of Medicine, National Institutes of Health, Bethesda, MD 20894, USA

## Abstract

In prokaryotic genomes, the number of genes that belong to distinct functional classes shows apparent universal scaling with the total number of genes [1–5] (Fig. 1). This scaling can be approximated with a power law, where the scaling power can be sublinear, near-linear or super-linear. Scaling laws are robust under various statistical tests [4], across different databases and for different gene classifications [1–5]. Several models aimed at explaining the observed scaling laws have been proposed, primarily, based on the specifics of the respective biological functions [1, 5–8]. However, a coherent theory to explain the emergence of scaling within the framework of population genetics is lacking. We employ a simple mathematical model for prokaryotic genome evolution [9] which, together with the analysis of 34 clusters of closely related microbial genomes [10], allows us to identify the underlying forces that dictate genome content evolution. In addition to the scaling of the number of genes in different functional classes, we explore gene contents divergence to characterize the evolutionary processes acting upon genomes [11]. We find that evolution of the gene content is dominated by two factors that are specific to a functional class, namely, selection landscape and genome plasticity. Selection landscape quantifies the fitness cost that is associated with deletion of a gene in a given functional class or the advantage of successful incorporation of an additional gene. Genome plasticity, that can be considered a measure of evolvability, reflects both the availability of the genes of a given functional class in the external gene pool that is accessible to the evolving microbial population, and the ability of microbial genomes to accommodate these genes. The selection landscape determines the gene loss rate, and genome plasticity is the principal determinant of the gene gain rate.

---

Power-laws are the simplest functions that give good fits to the data on gene scaling. However, given that genome sizes barely span two orders of magnitude (Fig. 1), these power functions should be treated as approximations rather than firmly established quantitative laws. These limitations notwithstanding, analysis of the scaling exponents using the power law approximation has shown that such exponents are (nearly) universal for each functional class across a broad range of microbes (notwithstanding some debate on the validity of the exact universality [4, 12]), suggesting that differences in scaling reflect important, not yet understood features of cellular organization and its evolution. In the seminal work on scaling, Van Nimwegen grouped the functional classes of genes along three integer exponents: 0,1,2, arguing that deviations from the integers most likely reflected gene classification ambiguities [5]. The gene classes with the 0 exponent include information processing systems (translation, basal transcription and replication), those with the exponent of 1 are primarily metabolic genes, and those with the exponent 2 are regulatory genes. In biological terms, the essential information processing systems are universally conserved and remain nearly the same in all microbes regardless of genome size; metabolic pathways expand proportionally to genome growth; and the complexity of regulatory circuits increases quadratically with the total number of genes. The toolbox model has been proposed to explain the quadratic scaling whereby the number of regulators grows faster than the number of metabolic enzymes thanks to the frequent re-use of the latter in new pathways [6, 7]. From the evolutionary standpoint, it has been suggested that the universal exponents are determined by distinct gene gain and loss rates for different classes of genes and represent the “innovation potential” of these classes [13]. Clearly, regulatory genes have the highest innovation potential whereas information processing systems have next to none. Here we formulate an explicit model for the gene gain and loss and express the scaling in terms of the evolutionary forces that emerge from our analysis, namely, the selection landscape and genome plasticity. The scaling that we obtain from this simple model does not follow a power law exactly but gives a comparable quality of fit within the range of available data.

**Figure 1.**
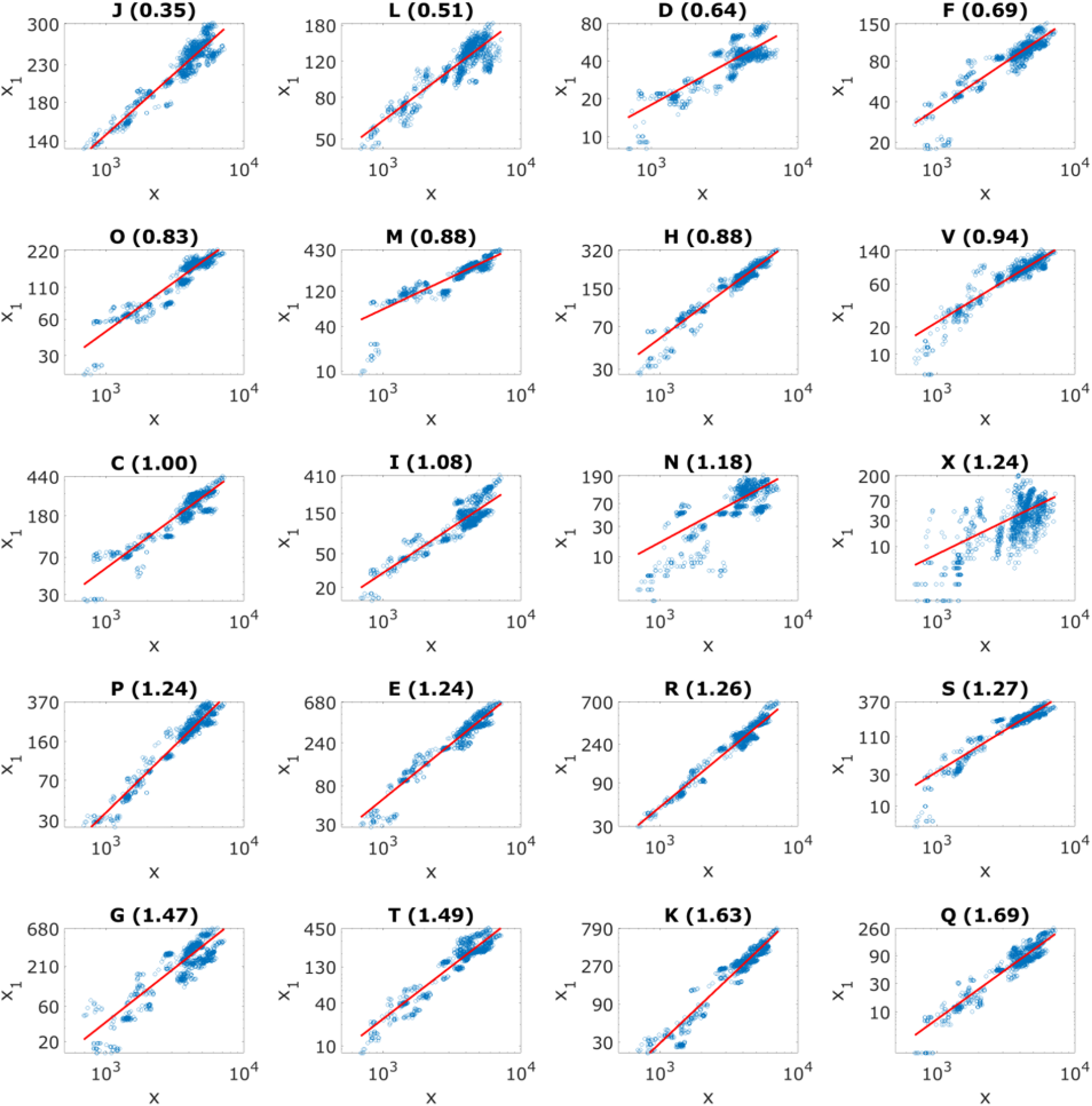
Scaling laws for all functional classes of the COGs. The number of genes in a given COG category is plotted against the total number of genes. Each point represents one genome from the analyzed set of 1490 genomes. The scaling is fitted to a power law which is indicated by a solid red line. The fitted scaling exponent is indicated in parentheses.

In the current study, we analyzed 20 functional classes of genes from the Clusters of Orthologous Groups (COGs) [14]. For the collection of microbial genomes analyzed here, the scaling exponents spun a range from 0.35 for translation genes (J COG category) to 1.69 for secondary biosynthesis genes (Q COG category) in our dataset (Table 1). It should be noted that the transcription category has an exponent of 1.63 because, in the COG classification, it includes both basal transcription proteins that, in the initial analysis, showed exponents close to 0, and transcription regulators, with the apparent quadratic dependence on the total number of genes. The analysis presented here imply that, in principle any scaling exponent is possible. Indeed, the observed values of gene category-specific exponents do not seem to perfectly fit the 0-1-2 paradigm but do show a broad range, with increasing exponents from the essential, universal information transmission genes to the more evolutionarily volatile genome components such as regulators and secondary metabolism enzymes. The robustness of the observed scaling exponents for different classes was tested by bootstrap analysis (Fig. S1; see Methods). Although, for some of the functional classes, the distribution of the bootstrap scaling exponents was wide ( *e.g.* secretion and motility genes (N); Fig. S1), the classes could be confidently partitioned into those scaling sub-linearly, near-linearly or super-linearly. The wide distributions also result in some pairs of classes overlapping (Table S1; see Methods). However, as shown below, similar scaling exponent can emerge from very different combinations of selection landscapes and genome plasticity ( *e.g.* secretion and motility genes (N) and the mobilome (X)).

**Table 1.** Scaling, selection and plasticity in different functional classes of microbial genes

| Class | Functions | scaling exponent | $\Delta S_1$ slope | average $\Delta S_1$ | average plasticity | plasticity slope |
| --- | --- | --- | --- | --- | --- | --- |
| J | translation | 0.35 | -1.10E-02 | 2.68 | 0.005 | 1.54E-05 |
| L | replication and repair | 0.51 | -1.35E-03 | 0.98 | 0.013 | -9.18E-05 |
| D | cell division | 0.64 | -1.58E-05 | 1.83 | 0.002 | -3.21E-05 |
| F | nucleotide metabolism and transport | 0.69 | -1.75E-02 | 2.15 | 0.003 | 3.65E-05 |
| O | posttranslational modification, protein turnover, and chaperone functions | 0.83 | -5.42E-03 | 1.38 | 0.010 | 4.88E-05 |
| M | membrane and cell wall structure and biogenesis | 0.88 | -1.05E-03 | 0.65 | 0.029 | 4.57E-05 |
| H | coenzyme metabolism | 0.88 | -4.68E-03 | 1.51 | 0.011 | 4.62E-05 |
| V | defense | 0.94 | -3.72E-07 | -0.44 | 0.034 | -6.68E-05 |
| C | energy production and conversion | 1.00 | -4.52E-03 | 1.19 | 0.017 | 8.92E-05 |
| I | lipid metabolism | 1.08 | -4.86E-03 | 0.92 | 0.016 | 1.24E-04 |
| N | secretion and motility | 1.18 | -7.13E-08 | 0.27 | 0.014 | 1.07E-04 |
| X | Mobilome: prophages, transposons | 1.24 | -2.06E-04 | -4.76 | 1.021 | 1.12E-02 |
| P | inorganic ion transport and metabolism | 1.24 | -3.76E-03 | 0.75 | 0.026 | 1.15E-04 |
| E | amino acid metabolism and transport | 1.24 | -1.90E-03 | 1.12 | 0.028 | 8.15E-05 |
| R | general functional prediction only | 1.26 | -1.05E-03 | 0.37 | 0.051 | 1.15E-04 |
| S | function unknown | 1.27 | -5.92E-07 | 0.55 | 0.025 | 3.35E-05 |
| G | carbohydrate metabolism and transport | 1.47 | -1.86E-03 | 0.46 | 0.041 | 1.65E-04 |
| T | signal transduction | 1.49 | -1.28E-03 | 0.57 | 0.030 | 1.25E-04 |
| K | transcription | 1.63 | -1.67E-03 | 0.27 | 0.058 | 1.81E-04 |
| Q | biosynthesis, transport, and catabolism of secondary metabolites | 1.69 | -5.77E-03 | 0.06 | 0.021 | 2.54E-04 |

We sought to uncover the evolutionary roots of the differential scaling of the functional classes of genes within the framework of the general theory of genome evolution by gene gain and loss. Prokaryotic genome evolution involves extensive horizontal gene transfer (HGT) and gene loss that can be expected to shape, among other features, the differential scaling [3, 15–17]. The simplest model for genome size dynamics describes the genomic evolutionary trajectory as a succession of stochastic gain and loss events [9]. The dynamics of the total number of genes in the genome *x* is therefore determined by the per genome gain and loss rates (*P*^+^ and *P*^−^), respectively

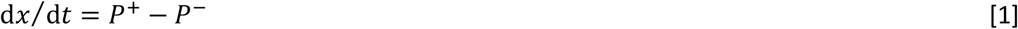

One of the key observable measures of microbial genome evolution is the pairwise intersection between genomes *I*, that is, the number of orthologous genes shared by a pair of genomes. Both the number of genes and the pairwise intersections between gene complements reflect genome content evolution and result from the same evolutionary processes. A complete theoretical description of genome evolution should therefore account for both these quantities. The stochastic gain and loss of genes entail a decay in pairwise genomes similarity through the course of evolution, even when the total number of genes remains approximately constant. As a first order approximation, pairwise genome intersections decay exponentially with the tree distance, with the decay constant *k* that is proportional to per-gene loss rate 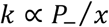 (see Methods for formal derivation). For an infinite gene pool [18]

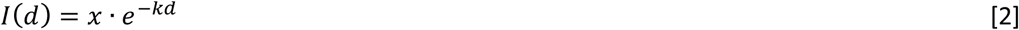

where *d* is the distance between the genomes along the tree. Given an infinite external gene pool, the rate of pairwise genome similarity decay is determined solely by gene loss rate. This model fits comparative genomic observations on the pairwise genome similarity decay with evolutionary distance in archaea, bacteria and bacteriophages [11] [19] [20]. We tested these observations on the ATGC set used for the present analysis and confirmed the close agreement of the model with the data (Fig. 2A, and Fig. S2).

**Figure 2.**
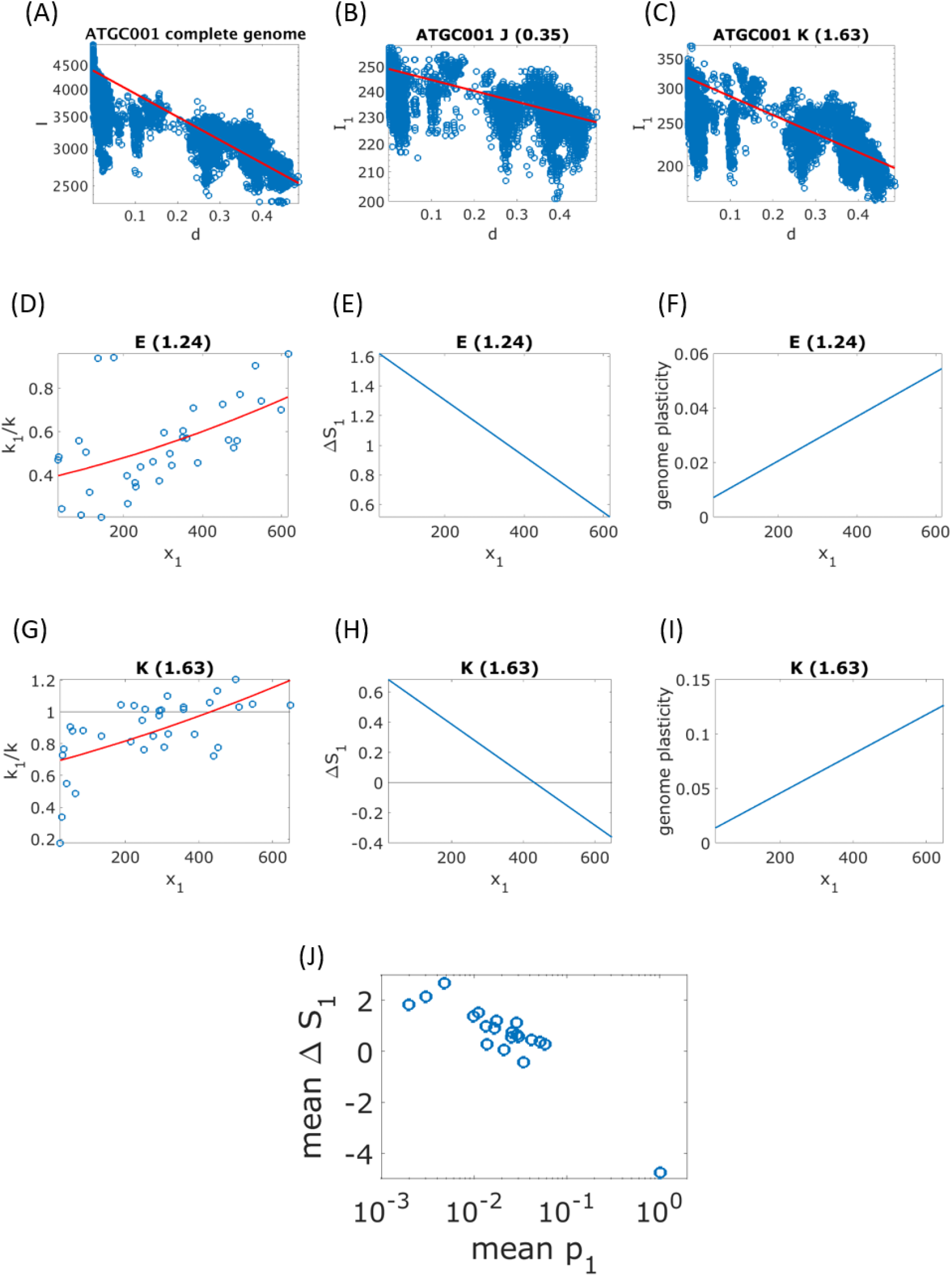
Decay of gene content similarity, selection landscapes and genome plasticity. **(A)** Pairwise genomes intersections plotted against tree distance *d* for complete genomes of ATGC001. Each point represents a pair of genomes in the ATGC, and the exponential decay fit of Eq. 2 is shown by the red solid line. **(B)** Pairwise genomes intersections for translation genes (J) from genomes of ATGC001. Each point represents a pair of genomes in the ATGC, and the exponential decay fit of Eq. 21 is shown by the red solid line. **(C)** Pairwise genomes intersections for transcription genes (K) from genomes of ATGC001. Each point represents a pair of genomes in the ATGC, and the exponential decay fit of Eq. 21 is shown by the red solid line). **(D)** Decay constant ratio *k*_1_/*k* is plotted against the number of genes in the functional category *x*_1_ for amino acid metabolism genes (E). Each point corresponds to an ATGC from the dataset. The model fit based on Eq. 11 together with the complete and class-specific selection landscapes of Eqs. 29 and 30, respectively, is shown by the solid red line. **(E)** The class-specific selection coefficient ∆*S*_1_ of Eq. 10 for amino acid metabolism genes (E), resulting from the fit shown in panel D. **(F)** Genome plasticity fitted using the linear approximation of Eq. 31 for amino acid metabolism genes (E). **(G)** Decay constant ratio *k*_1_/*k* is plotted against the number of genes in the functional category *x* _1_for transcription genes (**K**). Each point corresponds to an ATGC from the dataset. The model fit based on Eq. 11 together with the complete and class-specific selection landscapes of Eqs. 29 and 30, respectively, is shown by the solid red line. **(H)** The class-specific selection coefficient ∆*S*_1_ of Eq. 10 for transcription genes (E), resulting from the fit shown in panel G. **(I)** Genome plasticity fitted using the linear approximation of Eq. 31 for transcription genes (K). **(H)** Mean ∆*S*_1_ plotted against mean plasticity, for all functional classes. Mean values were calculated by averaging over all ATGCs.

To account for the dynamics of distinct functional classes of genes, we define gain and loss rates for the respective subsets of genes. Like the complete genome, each functional class (*x*_1_) is subject to stochastic gains and losses of genes that occur with rates 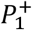 and 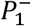, respectively

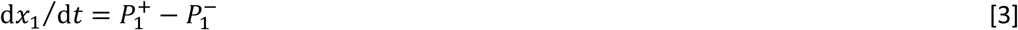

Below we express gain and loss rates explicitly and show how 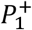 and 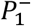 are related to the overall genome gain and loss rates *P*^+^ and *P*^−^. With respect to the genome content, all quantities can be defined for genomic subsets that include only genes from a specific functional class. We define class-specific pairwise intersection (i.e. the number of genes of class 1 shared between the pair of genomes) *I*_1_. Similar to its complete genome analog, the class-specific pairwise intersection decays exponentially with evolutionary distance. The decay constant *k*_1_ is proportional to the class-specific per-gene loss rate 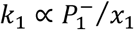. Empirically, gene classes with sublinear exponents are characterized by slow decay of pairwise intergenome similarity whereas those with super-linear exponents show fast decay (Figs. 2A-C and supplementary Figs. S3-S22).

Assuming finite effective population size with the weak genome dynamics limit (gain and loss rates are low enough such that gains and losses, hereafter “mutations”, occur and get fixed sequentially), gain and loss rates can be expressed as the product of the mutation rate and the probability for the mutation to get fixed in the population [9]. Mutation events are either an acquisition or a deletion of one gene, with the respective rates α and β. Accordingly, gain and loss rates can be written as

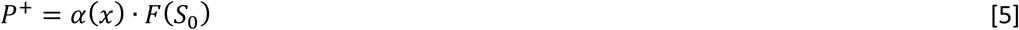

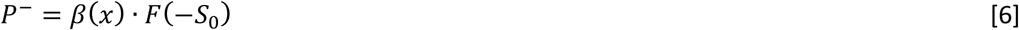

where *F* is the fixation probability and *S*_0_ is the genomic mean of the selection coefficient normalized by effective population size (see Methods). The *S*_0_ value can be regarded as the mean selective benefit (or cost) associated with the acquisition or loss of a random gene. Specifically, Eqs. 5 and 6 imply a symmetry in the selective effect with respect to gain and loss of a single gene: the benefit (or cost) is of equal magnitude for both events but with opposite signs [9, 21]. However, a closer examination of the gene acquisition process reveals a more complicated picture that involves two distinct time scales. Even genetic material that is beneficial on a large time scale, appears to be slightly deleterious initially, and fitness is recovered only after a transient time of several hundred generations [22]. In contrast, the coefficient *S*_0_ is inferred from extant genomes and thus reflects the average cost (or benefit) of gene deletion, and accordingly, the long-term average benefit (or cost) carried by a gene already incorporated in the genome. Within this formulation, the short time scale, that is, the transient phase of gene acquisition, is accounted for by the gain rate *α*. Specifically, *α* represents the product of the raw acquisition rate and gene acceptability, that is, the probability that the acquired gene is not rejected by the population within the short time scale.

Gain and loss rates for genes that belong to a specific functional class can be expressed following a similar reasoning. The class-specific selection landscape that determines the fixation probability term can differ from the mean selection landscape of the complete genome. We first develop the formulation of the loss rate which, under the assumption that deletions occur at random loci across the genome, is given by the complete genome deletion rate *β* multiplied by the fraction of the genome that is occupied by genes of a specific functional class. Together with the fixation probability for a deletion event that depends on the class-specific mean selection coefficient, *S*_1_, this gives

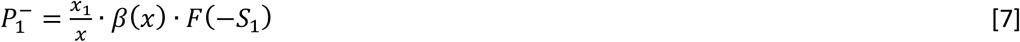

for the class-specific loss rate. The acquisition rate for class-specific genes is given by the product of the global acquisition rate *α*, fixation probability that depends on the class-specific mean selection coefficient, *S*_1_, and the class-specific genome plasticity *p*:

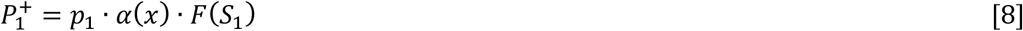

where the product *p*_1_. **α** denotes the probability that an acquired gene belongs to the specific functional class. As in the complete genome case, this formulation of class-specific gain and loss rates implies a symmetry between gain and loss, with respect to the selective effect. Accordingly, *S_1_* quantifies the long-term benefit or cost. If the short-term behavior is similar across all genes, the probability of a successful uptake of a gene is taken into account in the category-specific gain rate of Eq. 8 by *α*. In this case, *p*_1_ simply represent the class-specific genes availability, that is, the fraction of class-specific genes in the external gene pool. However, as described in detail below, the analysis of the scaling laws together with the pairwise intersection of the gene sets shows that *p*_1_ is genome size-dependent and does not fit the assumption of uniform acceptability across all classes of genes. The coefficient *p*_1_ therefore reflects not only the availability of class-specific genes, but also the class-specific ability of the microbial cell to tolerate additional genes of the given functional class within the short time scale. Hence we denote 1 class-specific genome plasticity.

Under the assumption that the genome size is approximately constant, the scaling laws can be derived from the relation between *x* and *x*_1_ that is expressed through the selection landscapes and genome plasticity (see Methods for derivation)

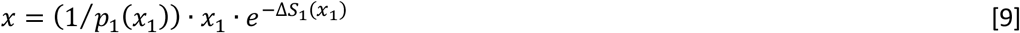

where ∆*S*_1_ is the mean selective (dis)advantage of a gene in the given functional class with respect to a random gene

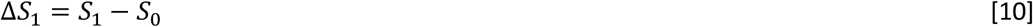

Eq. 9 describes the scaling of the number of genes in a functional class with the total genome size, and can be interpreted as follows. If class-specific genome plasticity *p*_1_ is independent of the number of genes in the class, the scaling is determined by ∆*S*_1_. For constant ∆*S*_1_, the scaling is linear, and the slope is greater (smaller) than *p*_1_ for genes that are on average more (less) beneficial than the genome-wide average, that is, ∆*S*_1_ > 0 (∆*S*_1_ < 0). Sublinear or super-linear scaling occurs for constant genome plasticity when ∆*S*_1_ depends on the number of genes ∆*S* _1_= ∆*S*_1_ (*x*_1_). Specifically, the scaling is sublinear (super-linear) when ∆*S*_1_ decreases (increases) with *x*_1_.

The derivation above provides the theoretical framework for inferring the class-specific selection landscapes and genome plasticity. The selection landscape determines the loss rate, whereas the genome plasticity is the principal determinant of the gain rate. The number of genes in a genome represents the balance between the two rates but pairwise genome intersections are determined by the loss rate alone. Thus, the genome intersection is a crucial ingredient in the analysis and allows us to disentangle selection landscape and genome plasticity, and determine the dependence of each of these factors on the number of genes. Because the scaling laws are robust with respect to local influences and are (nearly) universal across all prokaryotes (see Fig. 1), the evolutionary forces underlying scaling are likely to be universal to this extent as well. In particular, we assume that the functional class-specific selection landscapes and genome plasticity are similar for all genomes. Recently, however, we have shown that genome size evolution is subject to local effects and is governed by taxon-specific factors [21], in addition to the universal factors. To circumvent this taxon-specificity, represented here by the genome-wide acquisition and deletion rates *α* and *β*, we normalize the class-specific decay constant *k*_1_by the genomic mean decay constant *k*, for each ATGC separately. This normalization cancels out the ATGC-specific factors and allows us to infer the universal selection landscape and genome plasticity. We show that both factors depend on the genome size and thus contribute to the shaping of the genome content, and specifically, the scaling laws. Throughout the analysis we rely on our previous results [21] for the genome-wide selection landscape *S*_0_ (see Methods).

In the following, we show that the observed scaling exponents, together with the class-specific selection landscape that emerge from pairwise intersection, are consistent only with genome plasticity that depends on the number of genes. We first infer the selection landscape from the pairwise intersections. The class-specific ∆*S*_1_, is inferred from the ratio between the class-specific decay constant and the genomic mean (see Methods for derivation)

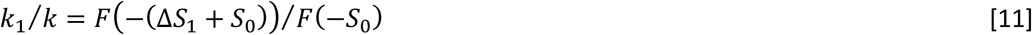

Given that we consider the ratio *k*_1_/*k*, the taxon-specific deletion rate *β* cancels out, and the ratio depends only on global factors, allowing an unbiased comparison among the ATGCs. The interpretation of Eq.11 is that genes that are associated with larger selection coefficients are exchanged less frequently than those that are subject to a weaker selection. For example, amino acid metabolism genes (E) show a *k*^1^/*k* ratio that increases with the number of genes (Fig. 2D), suggesting that the fitness cost of deletion of genes in this class drops for larger genomes. This behavior is typical and common to most functional classes, with the notable exception of defense genes (V) and the mobilome (X; the entirety of integrated mobile genetic elements) (Fig. S23). Accordingly, ∆*S*_1_ decreases with the class-specific number of genes *x*_1_(Fig. 2E and Fig. S24). However, as explained above, constant plasticity combined with ∆*S*_1_ that decreases with genome size, result in a sublinear scaling (see Eq. 9). The only way to reconcile the decreasing selection coefficient and super-linear scaling is to introduce genome size-dependent genome plasticity *p*_1_= *p*_1_(*x*_1_). The next step in the analysis is therefore to infer the genome plasticity, which can be extracted from the gain probabilities ratio (see Methods for derivation)

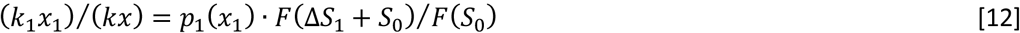

Similarly to Eq. 11, the genome-wide acquisition rate α, which can be subject to local influences [21], cancels out, allowing us to infer the selection landscape and genome plasticity from Eqs. 11 and 12. For simplicity, we use linear approximations for ∆*S*_1_(*x*_1_) and for *p*_1_(*x*_1_), to fit the data (Figs. 2E and 2F, and Supplementary Figs. S23 - S26; see Methods for details).

To better understand how the number of genes in each class is determined by the selection landscape and genome plasticity, it is useful to compare different classes in some detail. For example, for amino acid metabolism genes (E), the *k*_1_/*k* ratio is below unity (Fig. 2D), and accordingly, ∆*S*_1_ is positive even for larger genomes (Fig. 2E). For this gene class, plasticity increases with the genome size (Fig. 2F), leading to the observed moderate super-linear scaling, despite the decrease in ∆*S*_1_ with *x*_1_ (see Eq. 9). In contrast, the abundance of transcription genes (K), primarily, regulators, grows with the genome size such that the *k*_1_/*k* ratio becomes greater than unity (Fig. 2G) which correspond to ∆*S*_1_turning negative (Fig. 2H). The higher abundance and the super-linear scaling of transcription genes (K) is therefore attributed to the genome plasticity of this class, which is twice as high as that for amino acid metabolism genes (E) (Figs. 2F and 2I). This interplay between the selection landscape and genome plasticity is common for all gene classes, and consequently, there is a strong negative correlation between the mean values of ∆*S*_1_ and genome plasticity (Fig. 2J; Spearman correlation coefficient *ρ* = −0.79 (p-val < 10^−3^).

Finally, we tested the model consistency by reconstructing the scaling laws using the fitted selection landscapes and genome plasticity. Specifically, for each gene class, the fitted selection landscape and genome plasticity were substituted into Eq. 9, (Fig. 3A). For most classes, the fit quality of our model was comparable to albeit slightly worse than that of the power law fit (Table S2). The immediate source of errors in model fitting is the linear approximations for ∆*S*_1_ and for genome plasticity. Although not optimal, a linear approximation was applied to minimize the number of assumptions and parameters in the model, and can be regarded as a first order expansion of the actual functions. It should be noted that, unlike with the direct power law fit of *x*_1_ vs *x* data, the parameters for the model-derived scaling were inferred from the combination of the number of genes and pairwise similarity decay rates in ATGCs (Eqs. 11 and 12), that is, measurable quantities that characterize genome evolution. For all functional classes, with the exception of the defense systems (V) and the mobilome (X), the relative selection coefficient is positive and decreases with the genome size (Fig. 2E and Supplementary Fig. S24). For all except 3 functional classes (L, replication and repair; D, cell division; and V, defense), genome plasticity increases with the number of genes (Fig. 2F and Fig. S26), that is, the larger the genome, the higher the probability that an additional gene can be incorporated into the corresponding functional networks. Both the plasticity slope and the mean plasticity strongly, positively correlate with the scaling exponent, with respective Spearman correlation coefficients *ρ* = 0.81 (p-val < 10_−3_) and ρ= 0.74 (p-val < 10^−3^) (Fig. 3B).

**Figure 3.**
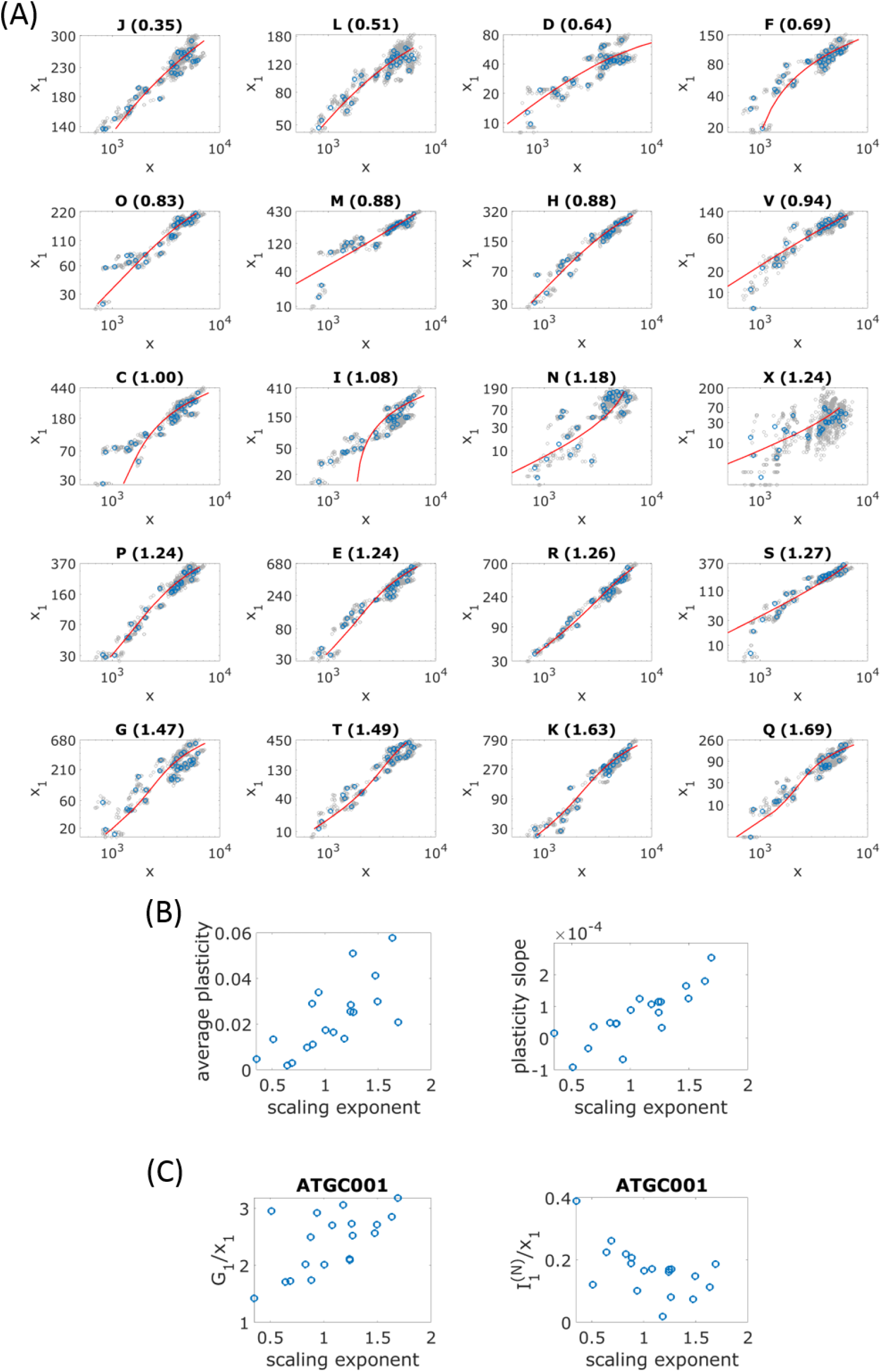
Model-derived scaling exponents for different functional classes of genes, genome plasticity, core genomes and pangenomes. **(A)** The number of genes in a COG functional category is plotted against the total number of genes. Blue points correspond to the mean values for each ATGC in the dataset. Individual genomes are indicated by gray points. The model fit of Eq. 9 is shown by the solid red line. **(B)** Average plasticity across all ATGCs and plasticity slope are plotted against the scaling exponent. Each point corresponds to a functional class of genes. The mobilome is associated with genome plasticity that is an order of magnitude greater than those of the other gene classes, and was excluded from the plot. **(C)** Class-specific pangenome G_1_ and core genome 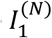 are plotted against the scaling exponent for ATGC001. Each point corresponds to a functional class of genes. To allow comparison between classes, pangenomes and core genomes are normalized by the number of genes in each class.

Functional classes with high plasticity, and accordingly, super-linear scaling exponents, are evolutionarily flexible and can be thought of as the microbial adaptation reserve. The biological properties of these classes appear compatible with this interpretation. Indeed, the 4 classes with the highest scaling exponents, namely, secondary metabolism (Q), transcription (K), signal transduction (T) and carbohydrate metabolism (G), are involved in reaction to rapidly changing environmental ques, including various biological conflicts (many of the Q category genes are involved in antibiotic production and resistance). These classes have high (G and K) or moderate (Q and T) plasticity and accordingly can accumulate in genomes to the point that the class-specific relative selection coefficient ∆*S*_1_ becomes negative so that these genes incur a non-negligible fitness cost on the organism. The genome similarity decay constant ratio *k*_1_/*k* for these functional categories is unity or greater in the majority of the ATGCs, that is, these genes are also lost at rates similar or higher than the average gene, resulting in their overall dynamic evolution. Notably, the gene categories with only a general functional prediction (R) and without any prediction (S) also showed super-linear scaling (albeit less pronounced than the above 4 classes) and high plasticity, suggesting that at least some of these genes contribute to adaptive processes. In agreement with previous results [23], we found that defense systems and the mobilome (the entirety of integrated mobile elements) incur a fitness cost on prokaryotes, and the relative cost of the mobile elements is an order of magnitude greater than that of defense systems. Not surprisingly, the genome plasticity of the mobilome also stands out, being at least an order of magnitude greater than that of all other classes (Table 1). Conversely, for sublinear classes, plasticity is low, so that incorporation of additional genes is unlikely albeit becoming more accessible in larger genomes. The genes in these classes are responsible for house-keeping functions that contribute less to short-term adaptation than the super-linear gene classes.

As a characteristic of the evolution of gene classes that can be directly determined from genome comparison, we analyzed the category-specific core genomes and pangenomes [24] (Fig. 3C). The normalized core genome and pangenome sizes correlate with the scaling exponent significantly and negatively for the core but positively for the pangenome, with the respective Spearman correlation coefficients *ρ* = −0.55 (p-val = 0.007) and *ρ* = 0.56 (p-val = 0.005). As expected, sublinear categories are associated with large relative core genomes and small relative pangenomes, compared to super-linear categories that make the principal contribution to the pangenome expansion. Thus, class-specific genome plasticity appears to shape the dynamics and architecture of microbial pangenomes.

To summarize, we provide here a general theoretical model explaining the universal scaling of the functional classes of genes in prokaryotes. The fits to the genomic data obtained with this model are comparable, even if slightly inferior to direct power law fits. This model does not include any assumptions on specific relationships between different functional classes as postulated in the previous models. Instead, we introduce an additional class-specific parameter that governs gene gain and loss processes, besides the selection coefficient, which we denote genome plasticity. Plasticity reflects the strength of purifying selection against horizontally acquired genes that has been previously described as the HGT barrier [25] as well as the availability of the genes of the given functional class which itself depends on their abundance in the external gene pool. Plasticity can be considered one of the forms of evolvability, a much debated concept [26–30] that, however, becomes the key factor shaping genome evolution in our model.

## Materials and Methods

### Genomic dataset

Clusters of closely related species from the ATGC database [10] that contain 10 or more genomes each were used in the analyses. The database includes fully annotated genomes and a phylogenetic tree for each cluster. Within each cluster of genomes, genes are grouped into clusters of orthologs (ATGC-COGs). Out of all genome clusters that contain 10 genomes or more, we selected the 36 genome clusters that match the following criteria: i) maximum pairwise tree distance is at least 0.1, and ii) the phylogenetic tree contains more than two clades, such that pairwise tree distances are centered around more than two typical values. Two of the 36 genome clusters were identified as outliers and were excluded from the dataset. The 34 genome clusters analyzed in this study are listed in Table S3. The ATGC-COGs were assigned to functional categories as defined in the COG database [14]. Genome sizes and sizes of functional classes of genes are given by the number of ATGC-COGs that are present in each genome and belong to the respective classes. Multiple genes from a single genome that belong to the same ATGC-COG were counted once. Genes without orthologs in other genomes (ORFans) genes were excluded from the analyses. Genome content analysis was performed for 20 COG categories. Functional classes of genes that were analyzed are listed in Table 1.

### Genome size evolution model

Substituting the gain and loss rates, *P*^+^ and *P^−^* of Eqs. 5 and 6, respectively, into the genome size dynamic of Eq. 1, we get the relation

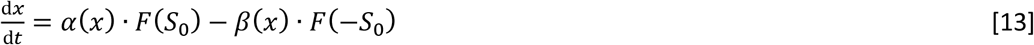

where scaling the time by the effective population size *N_e_*, allows to express gain and loss rates through *S*_0_ = *N_e_s_0_*, where *s*_0_ is the genome=wide average of the selection coefficient. Finally, we used the fact that, if an acquisition event is associated with selection coefficient *S*_0_, a deletion event would be associated with selection coefficient -*S*_0_ [9, 21]. The population size-scaled fixation probability *F* can be written as [31]

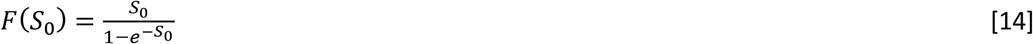

For a steady state, where *P*^+^ = *P*^−^, the selection and deletion bias are related by

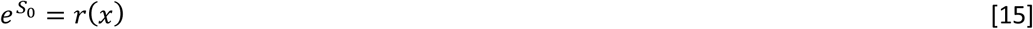

where the deletion bias is defined as r = *β*/*α*. The equation above reflects the selection-drift balance.

### Distinct functional classes of genes

In analogy to the stochastic equation for complete genome size dynamics, the dynamics of the number of genes that belong to a distinct functional class, denoted by *x*_1_, can be obtained by substituting the category-specific gain and loss rates of Eqs. 7 and 8, respectively, into Eq. 3

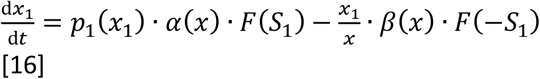

We assume a steady state and set d*x*_1_/d*t* = 0 in the equation above. Expressing the deletion bias *r* = *β*/*α* by the complete genome selection coefficient *S*_0_ using Eq. 15, we get the steady state relation of and *x*_1_, given by Eq. 9.

### Pairwise genome intersections *I*

To account for the genome content similarity, each genome is represented by a vector ***X*** with elements that assume values of 1 or 0. Each entry represents an ATGC-COG, where 1 or 0 indicate presence or absence, respectively, of that ATGC-COG in the genome. Genome size *x* is then given by the sum of all elements in ***X***. The number of common genes *I* is defined as

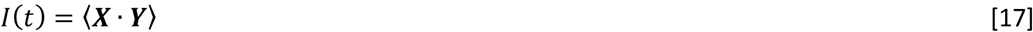

where ***X*** and ***Y*** are two vectors that represent the two genomes, the angled brackets indicate averaging over all possible pairs of genomes, and the dot operation stands for a scalar product. The pairwise genomes intersection dynamic is given by

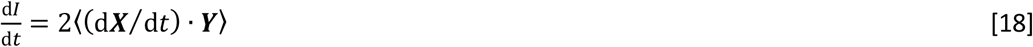

where we used the fact that both averages are equal 〈(d***X***/d*t*)∙ ***Y***〉 = 〈***X*** ∙ (d***Y***/d*t*)〉. Assuming a finite gene pool of size *L*, we have

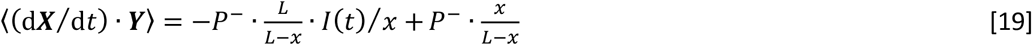

where the last approximation relies on the steady state assumption *P*^+^ ≈ *P*^−^. Substituting the relation above into the equation for the pairwise genome similarity time derivative and solving the differential equation, we obtain the exponential decay of the pairwise genome intersection to an asymptote *x*^2^⁄*L*

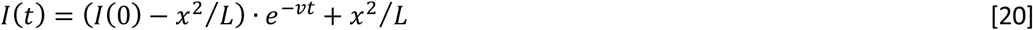

with decay constant

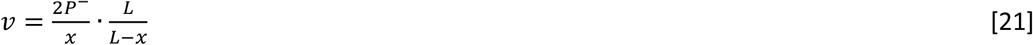

Assuming a clock with respect to loss events, the time *t* can be translated into tree pairwise distance as *d* = 2*t*⁄*t*_0_. Further assuming that the gene pool is much larger than the mean genome size *L* ≫ *x*, the pairwise similarity decays exponentially with respect to tree distance *d* as

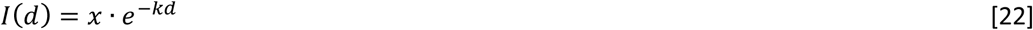

with decay constant

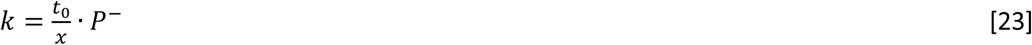

Note that the ratio *P*^−^⁄*x* gives the per-gene loss rate. It is possible to consider pairwise genome intersections with respect to a subset of genes. The derivation of Eqs. 17–23 can be repeated for genes that belong to a specific functional class. The functional class-specific genome intersection is therefore given by

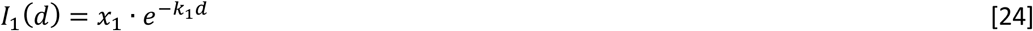

with decay constant

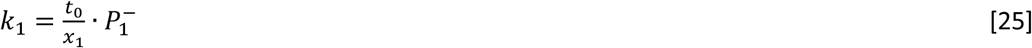

Note that for the ratio *k*_1_/*k*, the time scaling constant *t*_0_ cancels out, and we have

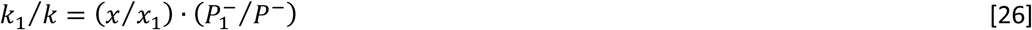

### Extraction of pairwise genome intersections decay constants from genomic data

Pairwise genome intersections *I* were calculated for all pairs of genomes in all genome clusters. Genome intersections were calculated for complete genomes as well as for different functional classes. Phylogenetic pairwise distances were extracted from the respective phylogenetic trees. The decay constants k and k_1_ were obtained by fitting the data to exponential decays (see below). Because ORFans genes were excluded from the dataset, the intercept was forced to the number of genes. Pairwise genome intersections are shown for all ATGCs for complete genomes and for all functional classes in Figs. S2-S22.

### Extraction of functional class-specific selection landscapes

To filter out taxon-specific factors [21] to the maximum extent possible, for each cluster of genomes we consider the category-specific quantities compared to the complete genome. Substituting the complete genome and class-specific gene loss rates of Eq. 6 and Eq. 8, respectively, into Eq. 26, we get the relation

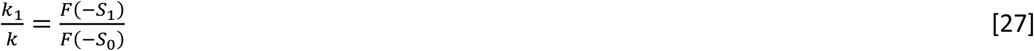

The class-specific selection landscape *S*_1_ is inferred from Eq. 27 as follows. The complete genome selection landscape *S*_0_ is known (see below), and the decay constants *k* and *k*_1_ are inferred from the data, as explained in the previous subsection. Finally, the genome plasticity is inferred using the gain rates. Under the steady state assumption, gain and loss rates are equal, such that Eq. 26 can be approximated by

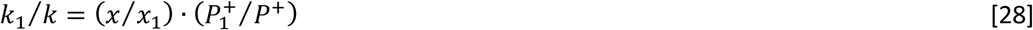

Substituting the complete genome and category-specific gain rates of Eq. 5 and Eq. 7, respectively, we get the equation for the genome plasticity

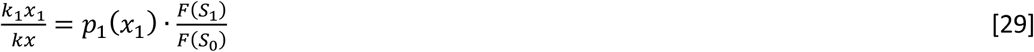

In a previous study, we found that the complete genome selection landscape *S*0 is related to the total number of genes by [21]

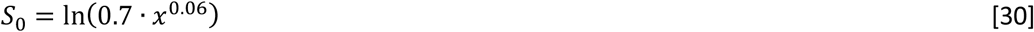

For simplicity, *S*_1_ is calculated relatively to *S*_0_, and the difference is taken to first order in *x*_1_

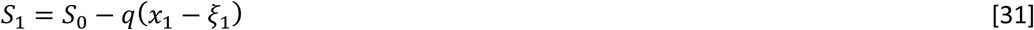

Similarly to ∆*S*_1_, the plasticity is taken as a first order function in *x*_1_

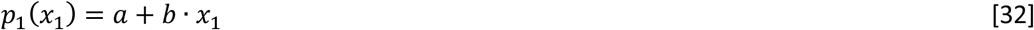

The resulting fits for the *k*_1_/*k* ratio of Eq. 26, and the ratio of Eq. 28 are shown for all COG categories in Figs. S23 and S25, respectively. Fitted relative selection landscape ∆*S*_1_ and genome plasticity are shown for all COG categories in Figs. S24 and S26, respectively.

### Data fitting and model parameters optimization

The numbers of genes in each class are discrete counts that typically span about one order of magnitude. Accordingly, it is assumed that the errors follow negative binomial distribution, and fitting was performed by optimizing model parameters together with the negative binomial distribution dispersion parameter, such that the log-likelihood is maximal.

### Inference of scaling power

Power law scaling exponents are obtained by fitting the genomic data to the curve *X*_1_ = *ƞ* ∙ *x*^γ^. For each functional class, parameters *a* and *γ* together with the negative binomial distribution dispersion parameter are optimized by maximizing the log likelihood for all genomes in the dataset. Genomes that do not contain genes that belong to the respective class were excluded from the analysis. The resulting fits are shown in Fig. 1, and the fit AIC values are listed in Table S2.

### Inference of pairwise intersection decay constants

The pairwise intersections decay constants *k* and *k*_1_ were inferred by fitting Eqs. 22 and 24 separately for each ATGC to the genomic data. The intercept is set to the mean number of genes (*x* for complete genomes and *x*_1_ for class-specific genes), such that the decay constant and the negative binomial dispersion parameter are optimized by maximizing the log-likelihood. Genomes that do not contain genes that belong to the respective class were excluded from the analysis. Fits are shown in Figs S2-S22.

### Optimizing model parameters

For each functional class, 4 model parameters, *q*, *ξ*_1_, *a* and *b* of Eqs. 31 and 32, are optimized using the mean numbers of genes and decay constants for each ATGC, *x*, *x*_1_, *k* and *k*_1_. Specifically, all 4 model parameters are optimized simultaneously using Eqs. 27 and 29, together with *S*_0_ of Eq. 30, by maximizing the goodness of fit *R*^2^ for both equations. Fits based on Eq. 27 are shown for all functional categories in Fig. S23, and those for Eq. 29 are shown in Fig. S25.

### Statistical analysis of scaling exponents

For each functional class, power law is fitted to a collection of genes generated by bootstrapping the original dataset. Specifically, the sampled dataset is generated by sampling with replacement the ATGCs, and collecting all genomes in sampled ATGCs. Sampling is performed over ATGCs and not directly at the level of genomes in order to avoid sampling bias due to the different number of genomes in each ATGC. The distribution of the fitted scaling exponents is shown for each class for 1000 bootstrap samplings in Fig. S1. For each pair of classes, the distribution overlap *C* is calculated. Specifically, for categories X and Y, with scaling exponents 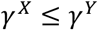 for the original dataset and bootstrap exponents 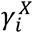and 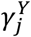, the overlap is given by

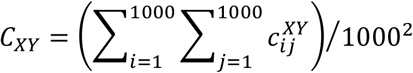

With

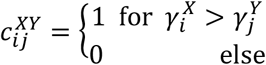

Given that, for the original dataset, the scaling exponent of class X is smaller than that of class Y, the overlap *C_XY_* indicates the probability of a bootstrap exponent of class X to be greater than the bootstrap exponent of class Y. Accordingly, *C_XX_* = 1⁄2.

## Supplementary figures and table captions

FIG. S1: Statistical support for scaling exponents calculated using bootstrap (see Methods). The distribution of fitted scaling exponents is shown for each class, for 1000 bootstrap samplings. The mean of the distributions is indicated by vertical dashed blue line, and the fitted scaling exponent for the original dataset is indicated by a vertical solid red line.

FIG. S2: Pairwise genome intersections I for complete genomes is plotted against tree distance *d*. Exponential decay fit of Eq. 2 is shown by a solid red line. The ATGC numbers are indicated in figure titles.

FIG. S3-S22: Pairwise intersections I1 for COG functional category J is plotted against tree distance d. Exponential decay fit of Eq. 21 is shown by a solid red line. The ATGC numbers are indicated in figure titles.

FIG. S23: Pairwise intersections decay constants ratio k1/k for all functional categories, together with fitted selection landscape (see Eq. 11). COG functional category name is indicated in the plot title, together with the functional category scaling exponent, which is indicated in parentheses.

FIG. S24: Fitted relative selection landscape ∆*S*_1_ for all functional categories (see Eq. 10). COG functional category name is indicated in the plot title, together with the functional category scaling exponent, which is indicated in parentheses.

FIG. S25: The ratio (k1×1) /(kx) for all functional categories, together with fitted selection landscape and genome plasticity (see Eq. 12). COG functional category name is indicated in the plot title, together with the functional category scaling exponent, which is indicated in parentheses.

FIG. S26: Fitted genome plasticity *p*(*x*_1_) for all functional categories. COG functional category name is indicated in the plot title, together with the functional category scaling exponent, which is indicated in parentheses.

TABLE S1: Overlap of scaling exponent bootstrap distribution of Fig. S1 (see Methods).

TABLE S2: Comparison of the fit quality to the genomic data for power law scaling and model-derived scaling (Eq. 9) in terms of Akaike Information Criterion (AIC).

TABLE S3: Genome clusters (ATGCs) in the analyzed dataset.

